# Integrative pipeline for profiling DNA copy number and inferring tumor phylogeny

**DOI:** 10.1101/195230

**Authors:** Eugene Urrutia, Hao Chen, Zilu Zhou, Nancy R Zhang, Yuchao Jiang

## Abstract

**Summary:** Copy number variation is an important and abundant source of variation in the human genome, which has been associated with a number of diseases, especially cancer. Massively parallel next-generation sequencing allows copy number profiling with fine resolution. Such efforts, however, have met with mixed successes, with setbacks arising partly from the lack of reliable analytical methods to meet the diverse and unique challenges arising from the myriad experimental designs and study goals in genetic studies. In cancer genomics, detection of somatic copy number changes and profiling of allele-specific copy number (ASCN) are complicated by experimental biases and artifacts as well as normal cell contamination and cancer subclone admixture. Furthermore, careful statistical modeling is warranted to reconstruct tumor phylogeny by both somatic ASCN changes and single nucleotide variants. Here we describe a flexible computational pipeline, MARATHON, which integrates multiple related statistical software for copy number profiling and downstream analyses in disease genetic studies.

**Availability and implementation:** MARATHON is publicly available at https://github.com/yuchaojiang/MARATHON.

**Supplementary information:** Supplementary data are available at *Bioinformatics* online.

## 1 Introduction

Copy number variations (CNVs) refer to duplications and deletions that lead to gains and losses of large genomic segments. CNV is pervasive in the human genome and plays a causal role in genetic diseases. With the dramatic growth of sequencing capacity and the accompanying drop in cost, massively parallel next-generation sequencing (NGS) offers appealing platforms for genome-wide CNV detection. In this note, we describe an analysis pipeline that integrates multiple aspects of CNV analysis, which can flexibly adapt to diverse study designs and research goals.

Despite the rapid technological development, CNV detection by high-throughput sequencing still faces analytical challenges due to the rampant biases and artifacts. Proper data normalization is crucial for sensitive and robust CNV detection, regardless of experimental designs and sequencing protocols. In whole-exome sequencing (WES) and targeted sequencing, where technical biases are usually magnitudes larger than CNV signals, data normalization is usually the pivotal step in affecting detection accuracy. Our proposed pipeline starts with data normalization using CODEX (Jiang, et al., 2015) and CODEX2 (Jiang, et al., 2017), which allow full-spectrum CNV profiling and are sensitive to both common and rare variants. Many large-scale genetic studies involve samples that have previously collected microarray data. It is currently unclear how microarray data can be used to improve sensitivity and robustness of the sequencing-based analyses. The pipeline proposed here seamlessly combine CODEX-normalized sequencing data with array-based log-ratio and B-allele-frequency measurements through iCNV (Zhou, et al., 2017).

Germline and somatic copy number changes are common in cancer and are associated with tumorigenesis and metastasis. In addition to the detection of total copy number changes, sequencing data give, at germline heterozygous loci, reads containing both alleles. This allows the disambiguation of allele-specific copy number (ASCN), which quantifies the number of somatic copies of each inherited allele. Compared to total copy number analysis, ASCN analysis gives a more complete picture of the copy number states, including copy-neutral loss of heterozygosity (LOH), which cannot be detected by total copy number analysis. Here we show how FALCON (Chen, et al., 2015) and FALCON-X (Chen, et al., 2017) integrate with CODEX and CODEX2 for estimation of ASCN.

Tumors are heterogeneous, genetically related populations of cells undergoing constant evolution. Much effort has been devoted to the reconstruction of the evolutionary phylogeny of tumors from bulk DNA sequencing data (Kuipers, et al., 2017). In addition to somatic ASCN changes, single nucleotide variants (SNVs) also provide valuable information for the reconstruction of the tumor phylogeny. We show that Canopy (Jiang, et al., 2016) for tracking of longitudinal and spatial clonal evolution can be applied to the outputs from FALCON and FALCON-X in an integrative analysis. This enables researchers to get both total and allele-specific DNA copy number calls and tumor phylogeny directly from BAM files.

## 2 Methods

The possible analysis scenarios are listed in Table 1. Figure 1 gives an outline for the relationship between the software: CODEX and CODEX2 perform read depth normalization for total copy number profiling; read depth normalized by CODEX/CODEX2 is received by iCNV, which combines it with allele-specific read counts and microarray data (if available) to detect CNVs; FALCON and FALCON-X perform ASCN analysis; and Canopy receives input from FALCON/FALCON-X to perform tumor phylogeny reconstruction. We propose MARATHON (copy nuMber vARiAtion and Tumor pHylOgeNy) as the integrated pipeline.

**Figure 1.**
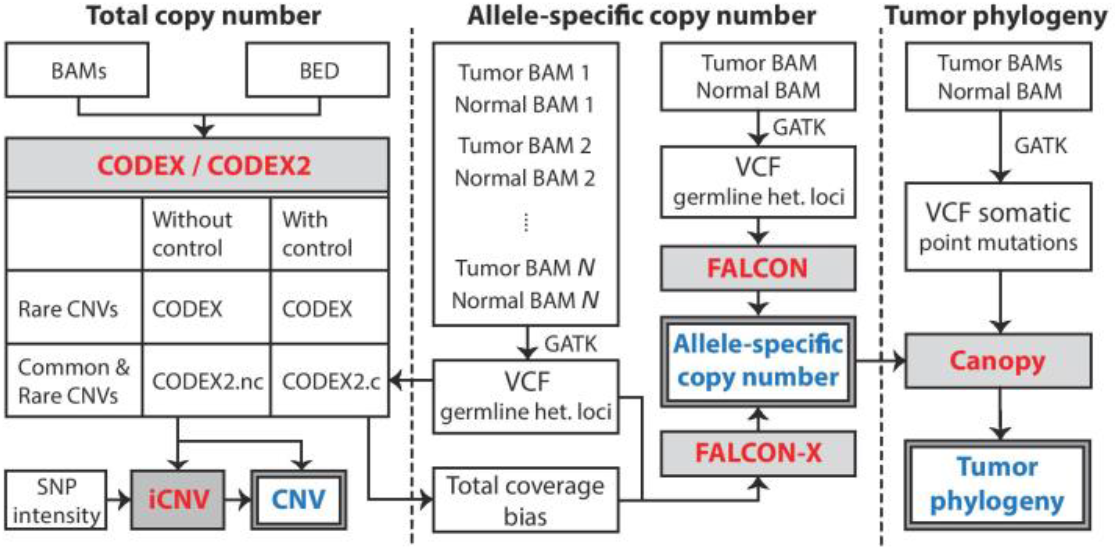
A flowchart outlining the procedures for profiling CNV, ASCN, and reconstructing tumor phylogeny. CNVs with common and rare population frequencies can be profiled by CODEX and CODEX2, with and without negative control samples. iCNV integrates sequencing and microarray data for CNV detection. ASCNs can be profiled by FALCON and FALCON-X using allelic read counts at germline heterozygous loci. Canopy infers tumor phylogeny using somatic SNVs and ASCNs.

**Table 1.** Analysis scenarios and pipeline design. The last column shows the sequence of software that should be used for each analysis scenario. *By “normal” we mean samples that are not derived from tumor tissue, which are not expected to carry chromosome-level copy number changes.

| Analysis Scenario |  |  | Pipeline |
| --- | --- | --- | --- |
| Goal | Study Design | Sample requirements |  |
| 1. Total copy number analysis of normal* | WES/WGS with or without matched array | >20 samples | CODEX2.nc → iCNV |
| 2. Total copy number analysis of tumor | WES/WGS /targeted sequencing | >20 normal controls (no need to be matched) | CODEX2.c |
| 3. Tumor allele-specific copy number | WGS | 1 tumor with matched normal sample | FALCON |
| 4. Tumor allele-specific copy number | WES/WGS | 1 or more tumor samples, multiple (>20) normal controls (no need to be matched) | CODEX2.c → FALCON-X |
| 5. Tumor phylogeny analysis | WGS | Multiple spatially or temporally separated tumor samples with matched normal sample | FALCON → CANOPY |
| 6. Tumor phylogeny analysis | WES/WGS | Multiple spatially or temporally separated tumor samples, multiple (>20) normal controls | CODEX2.c → FALCON-X → CANOPY |

CODEX (Jiang, et al., 2015) adopts a Poisson latent factor model for normalization to remove biases due to GC content, exon capture and amplification efficiency, and latent systemic artifacts. CODEX2 (Jiang, et al., 2017) builds on CODEX with a significant improvement of sensitivity for both rare and common variants. CODEX2 can be applied to two scenarios: the case control scenario (CODEX2.c in Figure 1) where the goal is to detect CNVs that are enriched in the case samples; and the scenario where control samples are not available (CODEX2.nc in Figure 1) and the goal is simply to profile all CNVs. CODEX and CODEX2 take as input assembled BAM files as well as bed files specifying targets for WES and targeted sequencing and output normalized read counts and tab-delimited text files with copy number calls. iCNV (Zhou, et al., 2017) uses the normalized coverage from CODEX/CODEX2, and makes use of sequenced reads at inherited single nucleotide polymorphism (SNP) positions for CNV detection. These heterozygous loci are shown to be valuable in improving detection and genotyping accuracy. If microarray data are available, iCNV also integrates log-ratios and B-allele frequencies from these platforms to boost accuracy and enable CNV detection in intronic regions. iCNV takes as input normalized coverage by CODEX/CODEX2, allelic frequency at inherited SNP positions from sequencing, and log-ratio and B-allele frequency from SNP array. Output is CNV calls with quality scores.

For ASCN estimation in a matched tumor-normal setting, FALCON (Chen, et al., 2015) is based on a change-point model on a process of a mixture of two bivariate Binomial distributions. FALCON takes as input allelic read counts at germline heterozygous loci by GATK (DePristo, et al., 2011) and outputs ASCN estimates with genome segmentations. For WES data, biases and artifacts cannot be fully captured by comparing the tumor sample to the matched normal sample. FALCON-X (Chen, et al., 2017) extends upon FALCON, where it takes as inputs allelic read counts at germline heterozygous loci and total coverage biases for each of these loci estimated by CONDEX2.c (Figure 1) and outputs ASCN estimates.

Canopy (Jiang, et al., 2016) identifies subclones within a tumor, determines the mutational profiles of these subclones, and infers the tumor’s phylogenetic history by NGS data from temporally and/or spatially separated tumor resections from the same patient. Canopy jointly models somatic copy number changes and SNVs in a similar fashion to non-negative matrix factorization and adopts a Bayesian framework to reconstruct phylogeny with posterior confidence assessment. Canopy takes as input both somatic ASCN changes returned by FALCON/FALCON-X as well as somatic SNVs and outputs tumor phylogenetic trees with somatic mutations placed along tree branches and subclones placed at the leaves.

## 3 Results

The proposed pipeline adapts to different study designs and research goals (Table 1). For population genetic and disease association studies, one would start with read depth normalization using CODEX/CODEX2, followed by CNV calling using iCNV. For cancer genomics studies where the goal is to obtain ASCNs and reconstruct tumor clonal history, one would start with read depth normalization using CODEX/CODEX2, followed by ASCN profiling using FALCON/FALCON-X and clonal history analysis using Canopy. R notebook with rich display is available for MARATHON. We also demonstrate in Supplementary Results a cancer phylogenetic study of a neuroblastoma patient (Eleveld, et al., 2015), as well as a breast cancer (Maxwell, et al., 2017) and a melanoma (Garman, et al., 2017) study where copy numbers are estimated.

## ACKNOWLEDGEMENTS

The authors thank Drs. Derek A. Oldridge, Sharon J. Diskin, and John M. Maris for providing the neuroblastoma dataset and Drs. Kara N. Maxwell and Katherine L. Nathanson for providing the breast cancer and melanoma dataset. *Funding:* This work was supported by the National Institutes of Health (NIH) grant P01 CA142538 to YJ and R01 HG006137 to NRZ. *Conflict of Interest:* none declared.

## References

Chen, H., et al. Allele-specific copy number profiling by next-generation DNA sequencing. Nucleic Acids Res 2015;43(4):e23.

Chen, H., et al. Allele-specific copy number estimation by whole exome sequencing. The Annals of Applied Statistics 2017;11(2): 1169–1192.

DePristo, M.A., et al. A framework for variation discovery and genotyping using next-generation DNA sequencing data. Nat Genet 2011;43(5):491–498.

Eleveld, T.F., et al. Relapsed neuroblastomas show frequent RAS-MAPK pathway mutations. Nat Genet 2015;47(8):864–871.

Garman, B., et al. Genetic and Genomic Characterization of 462 Melanoma Patient-Derived Xenografts, Tumor Biopsies, and Cell Lines. Cell Rep 2017;21(7):1936–1952.

Jiang, Y., et al. CODEX2: full-spectrum copy number variation detection by high-throughput DNA sequencing. bioRxiv 2017:211698.

Jiang, Y., et al. CODEX: a normalization and copy number variation detection method for whole exome sequencing. Nucleic Acids Res 2015;43(6):e39.

Jiang, Y., et al. Assessing intratumor heterogeneity and tracking longitudinal and spatial clonal evolutionary history by next-generation sequencing. Proc Natl Acad Sci U S A 2016;113(37):E5528–5537.

Kuipers, J., Jahn, K. and Beerenwinkel, N. Advances in understanding tumour evolution through single-cell sequencing. Biochim Biophys Acta 2017;1867(2):127–138.

Maxwell, K.N., et al. BRCA locus-specific loss of heterozygosity in germline BRCA1 and BRCA2 carriers. Nat Commun 2017;8(1):319.

Zhou, Z., et al. Integrative DNA copy number detection and genotyping from sequencing and array-based platforms. bioRxiv 2017:172700.

